# Rapid neural representations of personally relevant faces

**DOI:** 10.1101/585133

**Authors:** Mareike Bayer, Oksana Berhe, Isabel Dziobek, Tom Johnstone

**Affiliations:** Berlin School of Mind and Brain, Department of Psychology, Humboldt-Universität zu Berlin, Berlin, Germany; Department of Psychiatry and Psychotherapy, Central Institute of Mental Health, Medical Faculty Mannheim/Heidelberg University, Mannheim, Germany; Centre for Integrative Neuroscience and Neurodynamics, School of Psychology and Clinical Language Sciences, The University of Reading, RG6 6AH Reading, UK; School of Health Sciences, Swinburne University of Technology, 3184 Hawthorn, Australia

**Keywords:** EEG-fMRI, emotion, faces, personal relevance, representation

## Abstract

The faces of those most personally relevant to us are our primary source of social information, making their timely perception a priority. Recent research indicates that gender, age and identity of faces can be decoded from EEG/MEG data within 100ms. Yet the time course and neural circuitry involved in representing the personal relevance of faces remain unknown. We applied simultaneous EEG-fMRI to examine neural responses to emotional faces of female participants’ romantic partners, friends, and a stranger. Combining EEG and fMRI in cross-modal representational similarity analyses, we provide evidence that representations of personal relevance start prior to structural encoding at 100ms, with correlated representations in visual cortex, but also in prefrontal and midline regions involved in value representation, and monitoring and recall of self-relevant information. Our results add to an emerging body of research that suggests that models of face perception need to be updated to account for rapid detection of personal relevance in cortical circuitry beyond the core face processing network.

---

Faces are the most important social and emotional stimuli we encounter in daily life. They tell us whether our fellow humans are strangers, enemies, or loved ones, and how they are feeling. As face experts, we possess distinct neural pathways for processing transient (e.g. emotion) and stable (e.g. identity) aspects of faces (Bruce and Young 1986; Kanwisher 2000). Yet investigating the temporal dynamics and neural circuitry of human face perception has proved a considerable methodological challenge.

EEG and fMRI research suggests that structural face encoding and representations of identity first occur about 170ms after stimulus onset in the ventral visual stream, specifically the fusiform gyrus (Horovitz et al. 2004; Rossion and Jacques 2012). Recent more nuanced EEG & MEG analyses have suggested that face identity, emotion, age, and gender, might be represented within 100 ms after stimulus onset (Cauchoix et al. 2014; Nemrodov et al. 2016; Vida et al. 2017; Dima et al. 2018). Interestingly, a recent MEG study reported enhanced early (70-100ms) encoding of gender and identity in familiar faces (actors), with less robust encoding of gender and identity in unfamiliar faces (Dobs et al. 2019). These studies suggest that particularly relevant aspects of faces may influence processing *prior to structural encoding*, although the MEG and EEG analyses do not provide the spatial specificity to investigate the network underlying these effects. The processing of familiar faces has generally been linked to more robust face detection and recognition (Gobbini et al. 2013; Barragan-Jason et al. 2015; Ramon and Gobbini 2018), however, an additional advantage has been shown for personally relevant faces, which seem to be processed in a qualitatively different way than famous faces (for reviews, see Sugiura 2014; Ramon and Gobbini 2018; Kovács 2020). EEG findings suggest that the earliest effects of *personal* relevance of faces occur around 170 ms after stimulus onset, corresponding to structural encoding (Ramon and Gobbini 2018), though effects at higher-order processing stages (e.g. indexed by the P3 component) are more robust (e.g., Guerra et al. 2012).

Consistent with these results, fMRI results have shown that the processing of personally familiar faces additionally recruits brain regions outside of the visual processing stream (the so-called extended face network), which are involved in monitoring of self-relevant information (medial prefrontal cortex), episodic memories (precuneus, anterior temporal cortex), person knowledge (temporoparietal junction; TPJ), and emotion processing (amygdala, insula) (Haxby et al. 2000; Gobbini and Haxby 2007; Natu and O’Toole 2011; Visconti Di Oleggio Castello et al. 2017).

Despite considerable progress, it remains unclear how quickly the genuine personal relevance of faces, such as those of our friends and loved ones, is decoded, and the extent to which different parts of the ‘extended’ face processing network are engaged at early versus late processing of personal relevance. In this study, we addressed these questions through simultaneous recording of EEG and fMRI. We presented heterosexual females in a romantic relationship with pictures of their romantic partner, a close male friend, and a male stranger, displaying fearful, happy and neutral facial expressions. The face identities were chosen in order to allow for a more fine-grained investigation of personal relevance: While partner and friend both represent personally relevant, familiar faces (as compared to a stranger), there is also a distinction between the partner and both friend and stranger, concerning feelings of romantic love. Including emotional expressions also allowed us to explore the temporal course at which emotional expressions (fear, happiness and neutral) are represented (Dima et al. 2018; Muukkonen et al. 2020).

To identify the time course and brain regions involved in representing different aspects of personally relevant faces, we combined EEG and fMRI data using representational similarity analyses (RSA; Fig 1) (Kriegeskorte 2008; Cichy and Oliva 2020). Based on previous results, we expected the involvement of the extended face processing network in the processing of personal relevance, possibly starting at the stage of structural encoding at 170 ms.

**Fig. 1.**
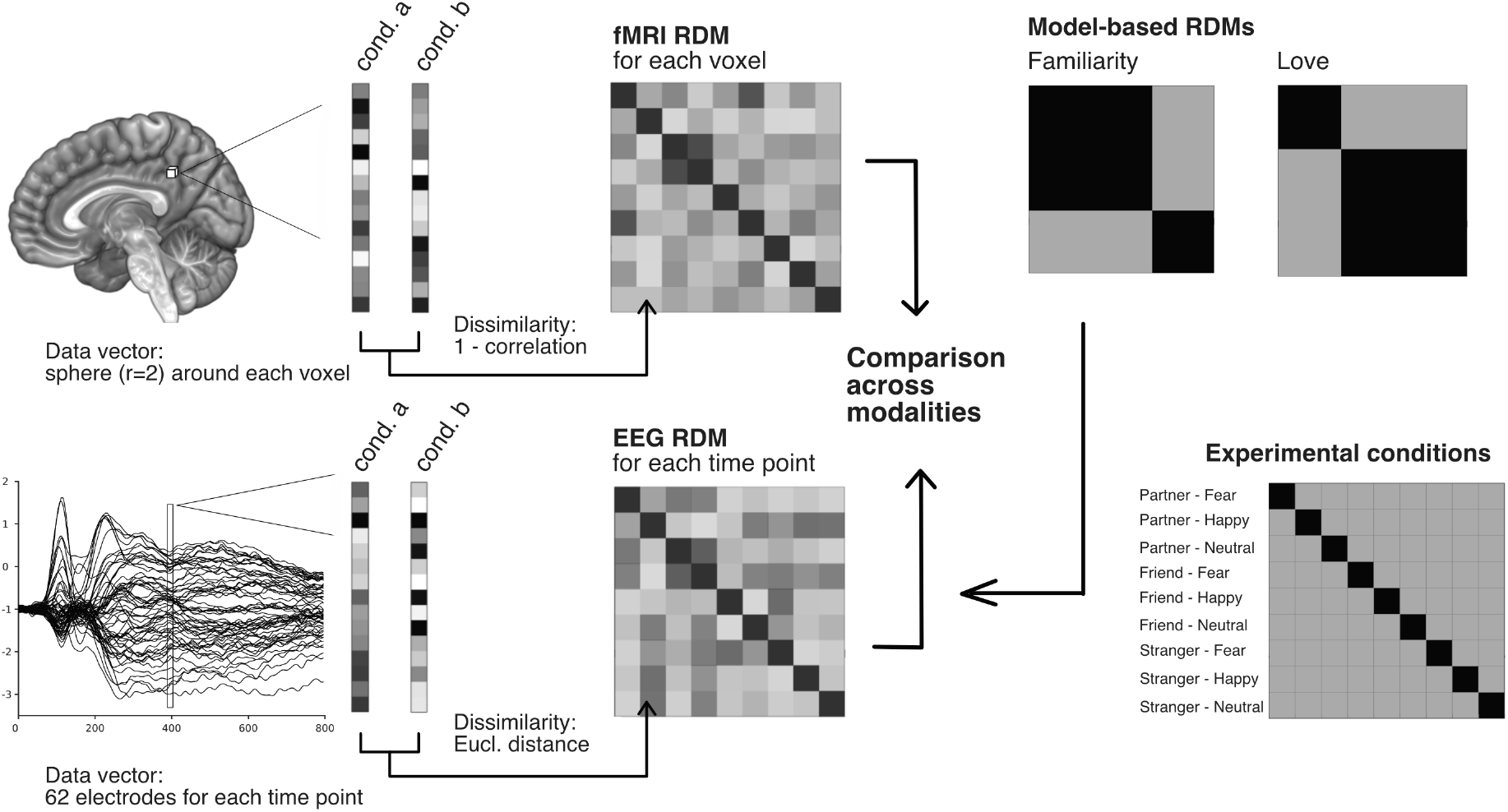
Representational Similarity Analyses. Representational dissimilarity matrices (RDMs) are constructed for each voxel in the brain (using a sphere around each voxel with a radius of 2 voxels) and for each data point in grand averaged ERPs by comparing pairwise, condition-specific activations. RDMs are symmetric with a diagonal of zeros, and their size corresponds to the number of experimental conditions, here 9 x 9. Model-based RDMs reflect theoretical predictions. After construction, RDMs are compared across modalities and with model-based RDMs.

## Materials and Methods

The study was reviewed and approved by the University of Reading research ethics committee. All participants provided informed consent before taking part in the study.

### Participants

Data were collected from 22 female participants (mean age = 19.8 years, sd = 0.9) recruited from the University of Reading undergraduate student population. All participants were in a heterosexual romantic relationship at the time of data collection (mean duration = 20.0 months, sd = 14.6; friendship: mean duration = 35.9 months, sd = 26.5). The decision to include only heterosexual female participants was based on experimental considerations, namely to limit our stimulus set to male faces in order to minimize differences in physical stimulus properties (including the use of a common stranger). Furthermore, that said group makes up the largest part of our recruitment population of psychology students. For four participants, no sufficient EEG data was available (for 2 subjects only 32 channel EEG was recorded, one participant was wearing a hair weave, one participant’s data was excluded due to excessive artefacts). Participants received a mean score of 105.9 points (sd = 10.7) of 135 points on the Passionate Love Scale (Hatfield and Sprecher 1986). They reported high contentment both with their relationship (mean = 8.8/10, sd = 1.3) and their friendship (mean = 8.3, sd = 1.3). All participants had normal or corrected-to-normal vision; 20 participants were right-handed. Participants were recruited through the Undergraduate Research Panel and Internet ads; they received course credit or £25 for participation.

### Stimuli

Stimuli consisted of portraits of the participant’s partner, a male close friend, and a male stranger, displaying fearful, happy, and neutral facial expressions (3 x 3 design). All stimuli were obtained by taking screen shots during a Skype session prior to the experimental session; all participants were presented with the same stranger. Stimuli were presented in color and displayed the face (including hair) of the depicted persons, who were directed to look straight at the camera. The background was removed, stimuli were presented on a grey background. For a representative example of stimuli, see Figure 2. After the main experiment, participants completed ratings of stimulus valence and arousal, as well as on attractiveness (neutral expression) using 7-point Likert scales. For rating values, see supplementary material.

**Fig. 2:**
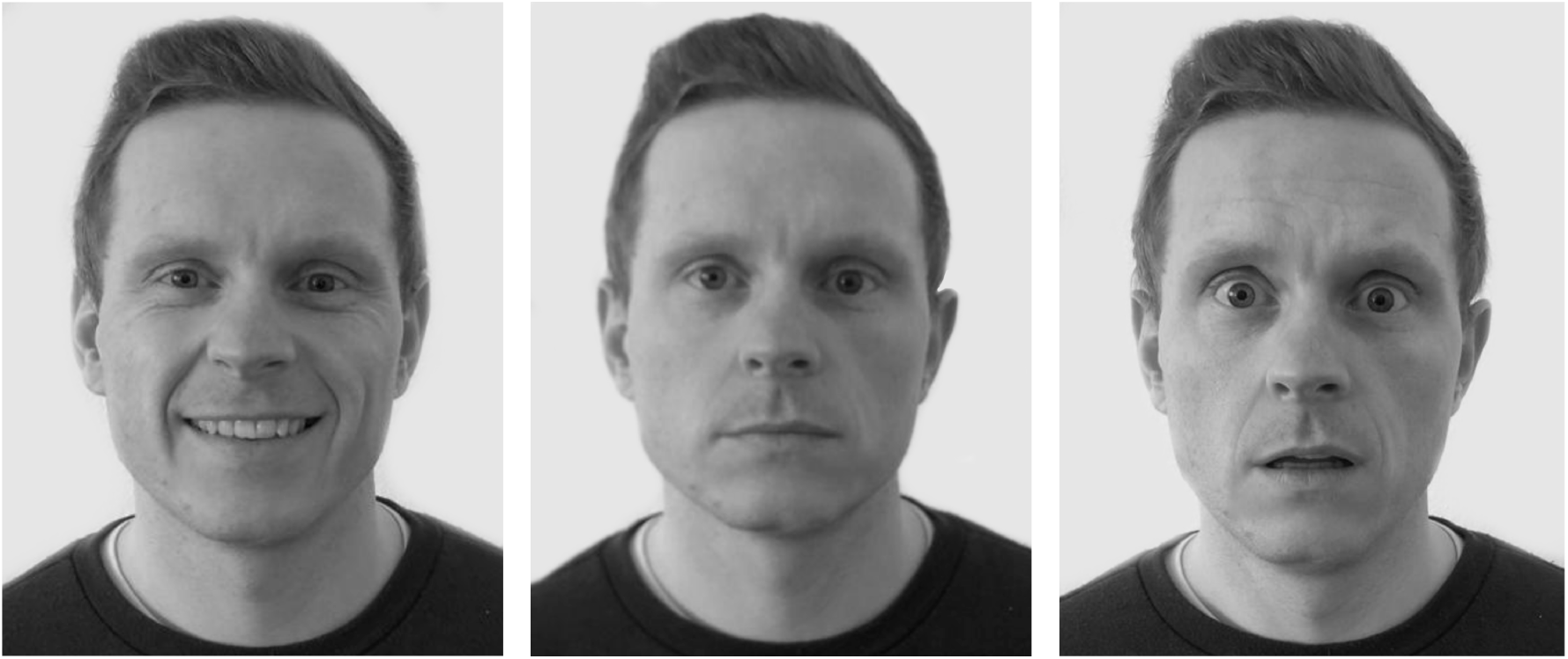
Example of experimental stimuli depicting a happy, neutral and fearful facial expression. In the original experiment, stimuli were depicted in color.

#### Procedure

After receiving information about the study, participants provided informed consent and were fitted with the EEG cap. Inside the scanner, participants performed a passive face-viewing task. Face stimuli were presented for 1s. Every face stimulus was presented 40 times, resulting in 360 experimental trials. Additionally, 40 1-back trials were presented to ensure participant’s attention to the faces. In these trials, participants had to indicate whether a face presented after a question mark was identical in identity and emotion to the one presented before the question mark. Stimuli were presented in 10 blocks. The stimulus sequence and the jittering of the inter-trial interval was determined using fMRI simulator (Rorden 2011) (mean ITI = 3000 ms, min = 2500 ms, exponential function). A central fixation cross was presented during the ITI. Stimuli were presented on a Nordic Neuro Labs goggle system at 60 Hz on an 800 x 600 pixel screen using EPrime software (Psychology Software Tools, Inc.).

### Data acquisition and pre-processing

#### fMRI

##### Acquisition

Data were collected on a 3T Siemens Trio MRI scanner using the standard 12 channel head coil. Functional images were acquired with a T2*-weighted gradient echo EPI sequence (40 interleaved axial slices, phase encoding P to A, 2.5 x 2.5 mm voxels, slice thickness = 2.5 mm, interslice gap = 0.25mm, 100×100 in-plane matrix, TR = 2500 ms, TE = 30 ms, Flip Angle: 90°). A high-resolution T1-weighted whole-brain structural image was acquired using an MPRAGE sequence (1mm isotropic voxels, FOV = 160 x 256 x 256 mm, Flip Angle: 9°).

##### Preprocessing and GLM

Data were processed with FSL 5.0 (http://www.fmrib.ox.ac.uk/fsl). Brain extraction was performed using the BET (Smith 2002). Data were denoised (Marchenko-Pastur Principal Component Analyses), motion corrected (Jenkinson et al. 2002), spatially smoothed using a 5 mm FWHM Gaussian kernel (Smith and Brady 1997), and grand-mean intensity normalized. Motion artefacts were removed with ICA-AROMA (Pruim et al. 2015) and data were high-pass filtered (100 s). Single subject 4D data were registered to the subject’s structural image using BBR (Jenkinson and Smith 2001). Registration of the individual structural image to the 2mm MNI 152 template was performed using ANTS (Avants et al. 2009). Finally, transformations were combined for the registration of the functional data to standard space. A first level GLM was applied with regressors for emotion by identity conditions (9 regressors) and for 1-back trials, created by convolving the temporal profile of each experimental condition with the FSL double gamma haemodynamic response function. Nuisance regressors without convolution were included to model breaks between blocks and artefacts (framewise displacement > 1mm).

#### RDMs

Representational dissimilarity matrices (RDMs; Kriegeskorte 2008; Cichy et al. 2016) were constructed for each voxel in the brain (more specifically, for a sphere around each voxel, see below), separately for each subject, based on z-statistic images of each of the 9 experimental conditions included in the mixed-effects GLM. For each voxel and condition, we created a vector (based on a sphere around each voxel, radius = 2 voxels). We quantified the dissimilarity between each pair of experimental conditions as 1 – Pearson’s R of the two corresponding vectors. This resulted in a 9 x 9 RDM for each voxel and each participant. For unimodal second-level fMRI analyses and results, see supplementary material.

#### EEG

##### Data acquisition

Continuous EEG data were collected from 64 Ag-AgCl electrodes simultaneously to fMRI acquisition (BrainProducts system); the electrocardiogram (ECG) was recorded from an electrode placed left of the spinal column. The sampling rate was 5000 Hz; data were referenced online to electrode FCz with electrode AFz as ground. Electrode impedances were kept below 20 kΩ. A sync box (BrainProducts) synchronised the MRI and EEG computer clocks.

##### Preprocessing

Offline, MR gradient artefacts were removed from continuous, baseline corrected data (using the whole artefact for baseline correction) with a sliding window of 21 artefacts (Allen et al. 2000). Data were low-pass filtered using a FIR filter (70 Hz) and down-sampled to 250 Hz. Ballistocardiographic artefacts were identified using the ECG with a semiautomatic template matching procedure and corrected using a template subtraction approach (sliding window of 15 pulse intervals). A restricted Infomax ICA (Bell and Sejnowski 1995) was used to remove eye blinks, eye movements and residual ballistocardiographic artefacts. Data was re-referenced to average reference and segmented into epochs from -100 ms to 800 ms relative to stimulus onset, and baseline-corrected using a 100 ms pre-stimulus baseline. Trials with activity exceeding ± 100 μV or voltage steps larger than 100 μV were excluded from analyses (0.6 % of trials). Data were averaged per participant and experimental condition.

#### RDMs

RDM matrices were constructed using across-subject grand-averaged waveforms, in order to minimize idiosyncratic sensory responses and to improve the signal-to-noise ratio (e.g., Cichy et al. 2016). For each time point from stimulus onset to 800 ms after stimulus onset, the distance between pairs of experimental conditions was quantified as their Euclidean distance across all 62 scalp electrodes. Euclidean distance was used in order to account for amplitude differences, which convey essential information in event-related potentials. Analyses resulted in a 9 x 9 RDM for each time point.

In order to constrain target EEG RDMs in the cross-modal analyses, we derived a measure of internal structure of the EEG RDM at each time point, in order to identify time points with potentially high representational content. For each time point’s RDM, we computed Euclidean distances of each RDM cell value to the arithmetic mean of all cell values in that RDM. As a result, EEG time points with pronounced differences between specific condition pairs relative to the mean difference between all pairs receive high values, whereas relatively smaller differences result in low values. For these calculations, we used one half of the (symmetrical) RDM, excluding the diagonal zeros. Time points of interest (peaks in EEG RDM structure) were identified at 52 ms, 108 ms, 204 ms, 308 ms, 428 ms, and 660 ms (see Fig. 4).

### Data analyses

#### Model-based EEG regression analyses

To characterize the information space represented by EEG-RDMs, we performed linear regression analyses on EEG RDMs using model RDMs as explanatory variables. In other words, we wanted to determine how experimental factors were represented throughout the EEG time line. Model RDMs were created based on the assumption of high similarity within and low similarity across conditions (coded as 0 and 1, respectively). We computed model RDMs for face familiarity (Partner/Friend vs. Stranger) and romantic love (Partner vs. Friend/Stranger). Emotion conditions were represented as emotional valence (distance between happy/fearful vs. neutral = 1, distance happy vs. fearful = 2) and arousal (distance happy/fearful vs. neutral = 1, distance happy vs. fearful = 0). For each EEG time point of interest (as revealed by the EEG structure measure), we entered RDMs for Familiarity, Love, Valence and Arousal as explanatory variables in the regression analysis. Additionally, in order to control for possible effects of low-level visual features, we constructed RDMs for each stimulus based on a model of cortical responses of neurons in the primary visual cortex (HMAX; Serre et al. 2007), more specifically on the output of the second layer (C1) of the model. RDMs for each participant’s individual stimulus set were calculated using Pearson’s correlations. For the EEG linear regression analyses, HMAX responses were averaged across participants with available EEG data. In addition to analyzing the time points of interest, we performed identical exploratory analyses for the complete time line from stimulus onset to 800 ms, and identified significant time windows (corrected p < .05) using a cluster-based permutation approach with a cluster-defining threshold of .05 and 1000 permutations.

#### EEG-fMRI fusion

After RDMs had been constructed for each voxel in the brain, EEG RDMs at time points of interest (as identified by our measure of internal structure) were used as target RDMs in single subject cross-modal searchlight analyses (using Pearson’s R), in order to identify brain regions with corresponding fMRI-derived representations. The resulting single subject RSA brain maps were entered into whole-brain second level analyses using permutation tests (1-sample t-test using FSL randomize, with 5000 permutations and an MNI gray matter mask with tissue priors thresholded at 0.3 and variance smoothing of 2mm). Threshold-free cluster enhancement was used to ensure a corrected Familywise Error (FWE) of < .05. Using permutation tests for second level analyses mitigates against bias in the first level RSA, since the permutation procedure produces a null distribution that incorporates the bias (hence if lower level estimates are positively biased, the null distribution is as well, and hence the criterion for significance at a given alpha is adjusted upwards).

#### EEG-fMRI regression analyses including visual feature models

In order to control for possible effects of low-level visual features on RSA results, we performed posthoc cross-modal RSA regression analyses that included individual HMAX RDMs (see above) as covariates. With these analyses, we aimed to assess whether results remained significant when accounting for representations of low-level stimulus features.

RSA analyses were performed using the CoSMoMVPA toolbox (Oosterhof et al. 2016) and the toolbox for representational similarity analyses (Nili et al. 2014) on Matlab R2017b.

## Results

Peaks in EEG RDM structure were evident at 52 ms, 108 ms, 204 ms, 308 ms, 428 ms, and 660 ms. Accordingly, these RDMs were used as searchlight RDMs in the whole-brain analyses on single-subject fMRI data (Fig. 4). Model-based EEG regression analyses of these time points showed significant correspondence of EEG representations with RDMs of Familiarity at 108 ms, 204 ms, 428 ms and 660 ms, and of Love at 308 ms, 328 ms and at 660 ms (see Fig. 3). Interestingly, none of the time points showed significant correspondence to RDMs of Valence, Arousal, or HMAX. In exploratory continuous analyses corrected by cluster-based permutation tests, significant correspondence of Familiarity was evident from 176 ms until 800 ms; and for the Love-RDM from 240 ms to 776 ms (see Fig. 3). Again, Valence, Arousal and HMAX did not show significant correspondence at any time point.

**Fig. 3:**
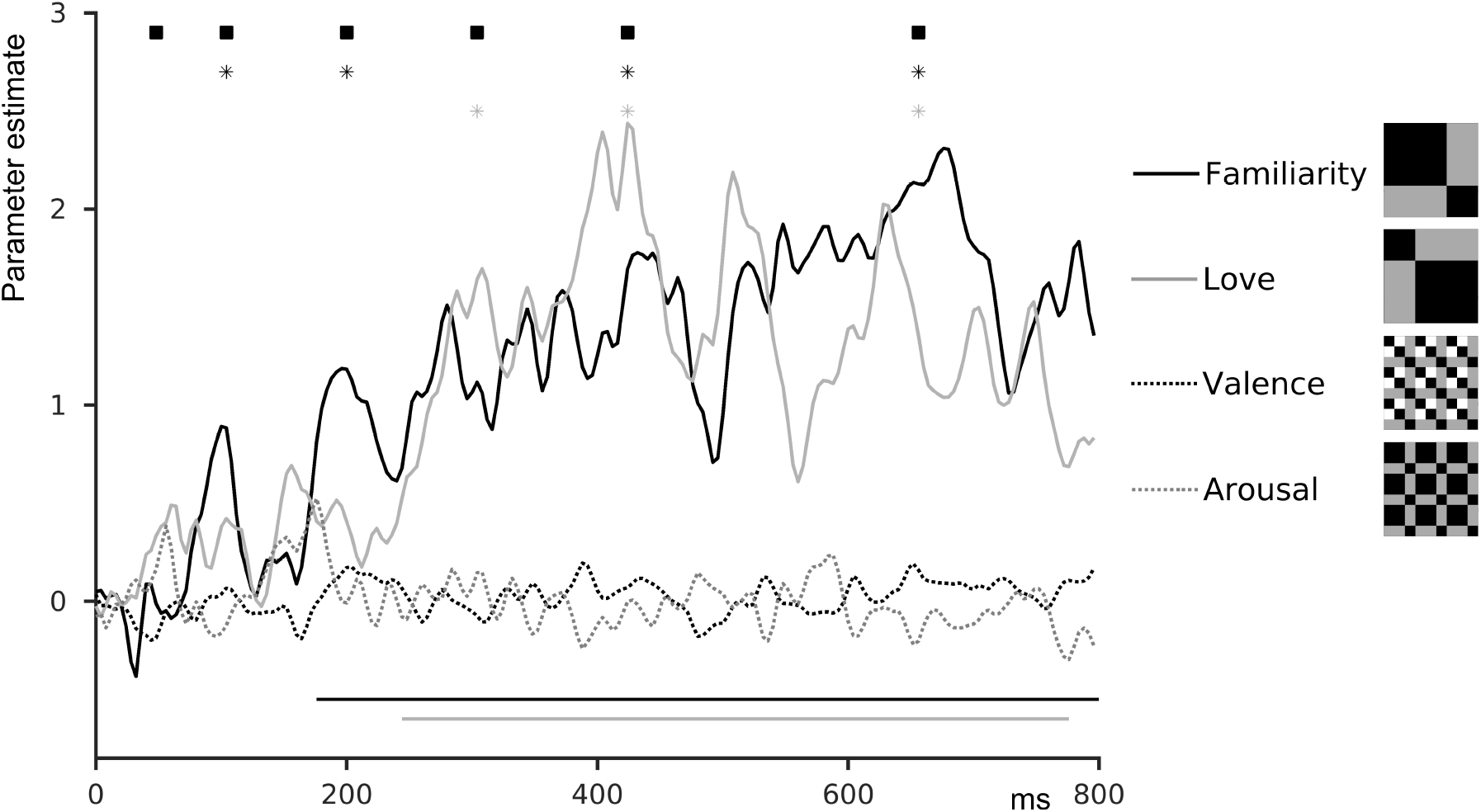
Parameter estimates for theoretical RDMs for Familiarity, Love, Valence and Arousal in linear regression analyses on EEG RDMs. Squares indicated time points of interest in the EG structure; stars mark significant activations (*p* < 0.01, uncorrected) for Familiarity and Love. Lines indicate significant time windows in continuous analyses determined by cluster-based permutation tests (corrected *p* < .05). Model RDMs for each variable are shown on the right.

**Fig. 4:**
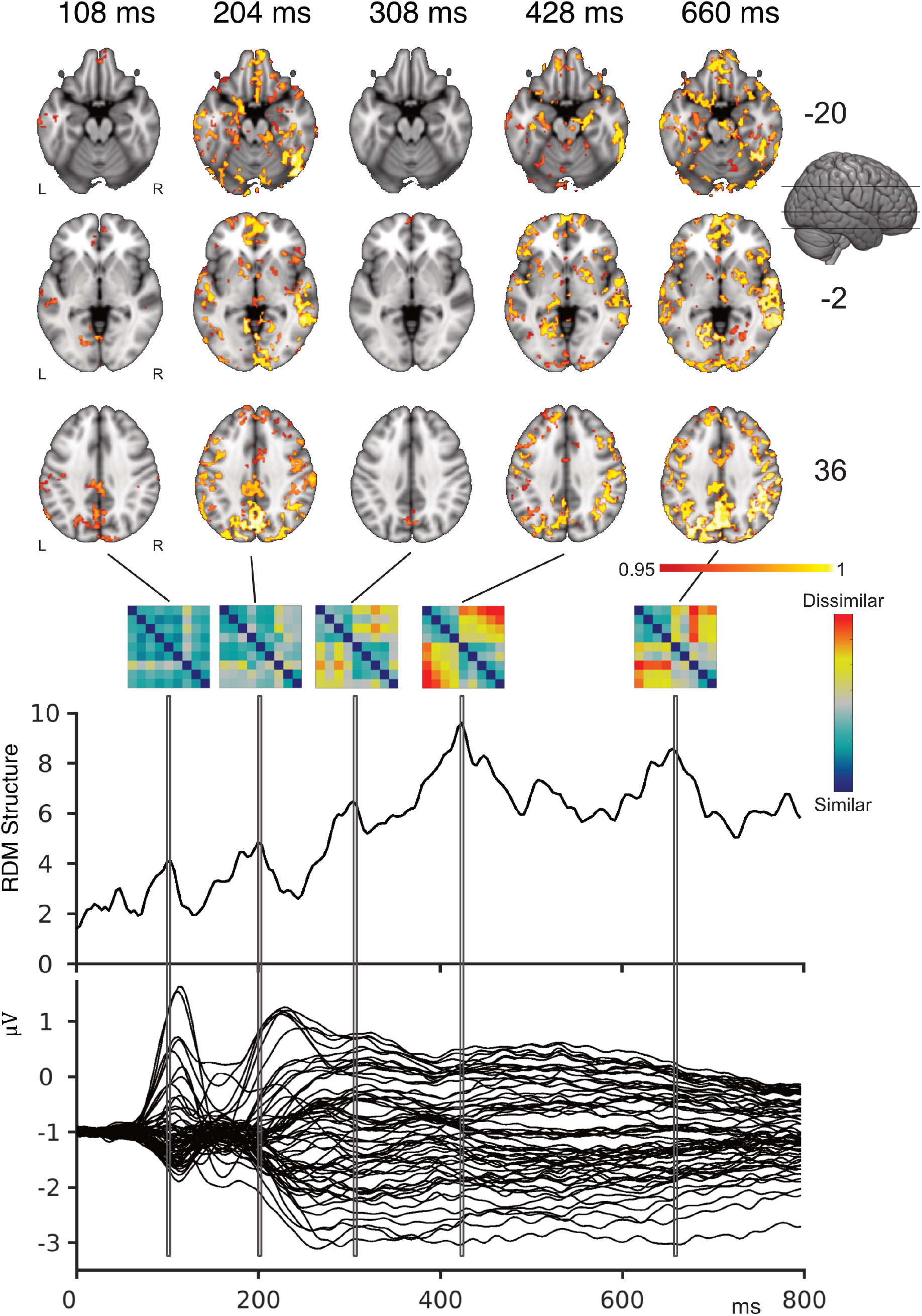
Combined EEG-fMRI representations. Brain regions in which fMRI RDMs corresponded to representations in the EEG at the corresponding time points. RDMs show EEG representations used as target RDMs in the combined EEG-fMRI analyses, selected on the basis of peaks in the RDM structure (top graph). Each line in the lower graph represents the mean time series for one electrode.

In combined EEG-fMRI analyses, there were no brain regions with significantly correlated spatial RDMs for the EEG RDM at 52 ms after stimulus onset. EEG representations at 108 ms showed correspondence to BOLD representations in the visual cortex (intracalcarine cortex and lingual gyrus), but also in the extended face network, including the the ventromedial PFC, precuneus, posterior cingulate cortex and TPJ. At 204 ms, more widespread corresponding EEG-fMRI representations additionally included the ACC, fusiform gyrus, amygdala, insula and N. accumbens. At 308 ms, representations were more focal and most evident in the precuneus, posterior cingulate and superior frontal gyrus. Finally, EEG-fMRI representations at 428 ms and 660 ms after stimulus onset were again more extended, with additional significant representations in the middle temporal gyrus, TPJ, in the inferior and superior frontal gyri, as well as in regions of the midbrain. Results of EEG-fMRI RSA analyses are depicted in Fig. 4.

Results of RSA regression analyses which included an RDM model of cortical responses in the visual cortex (HMAX) confirmed our previous analyses, showing that significant representations were not determined by lower-level stimulus features. See Table 1 for a complete list of brain regions and the results of RSA HMAX regressor analyses.

**Table 1.**
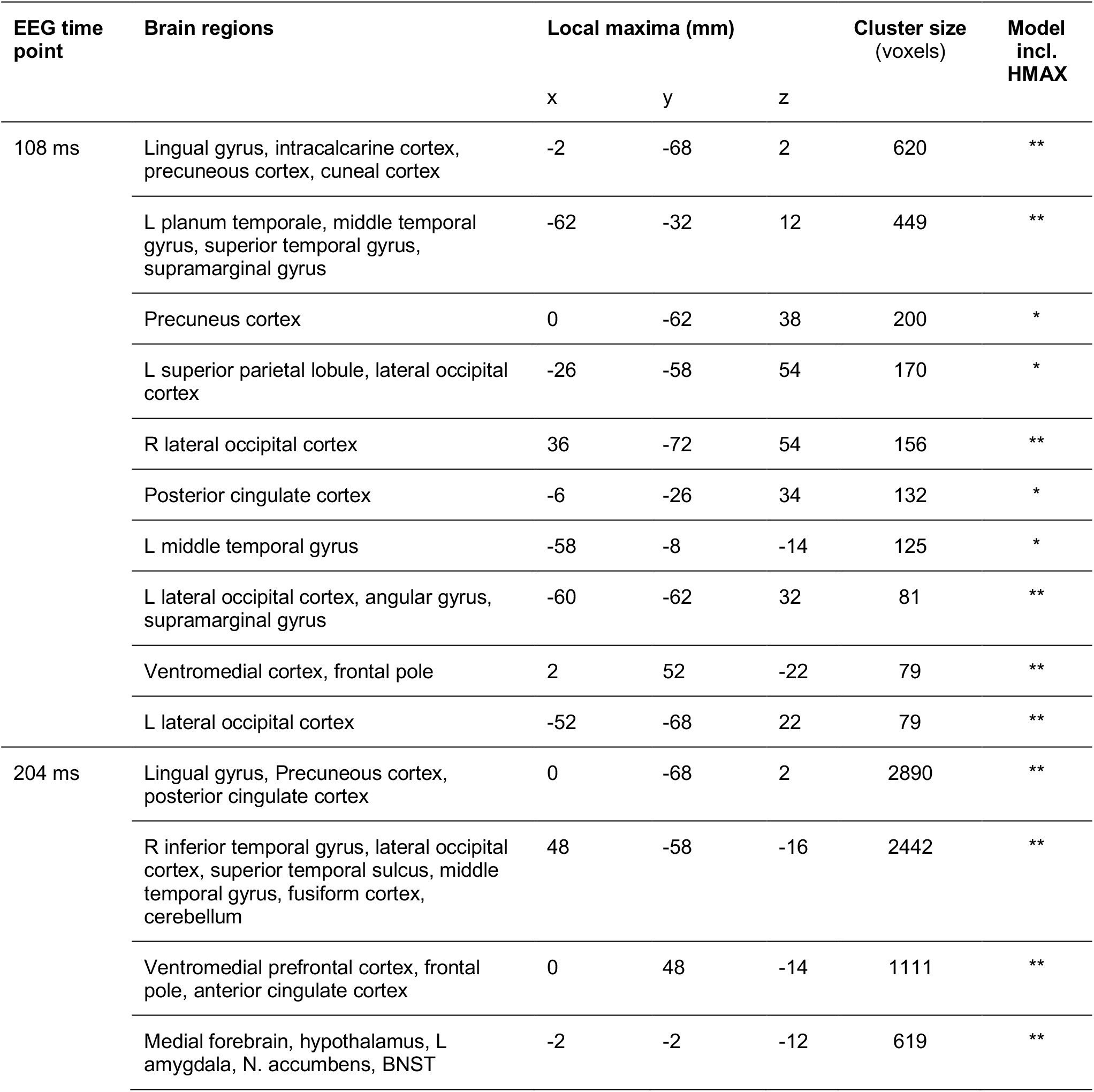
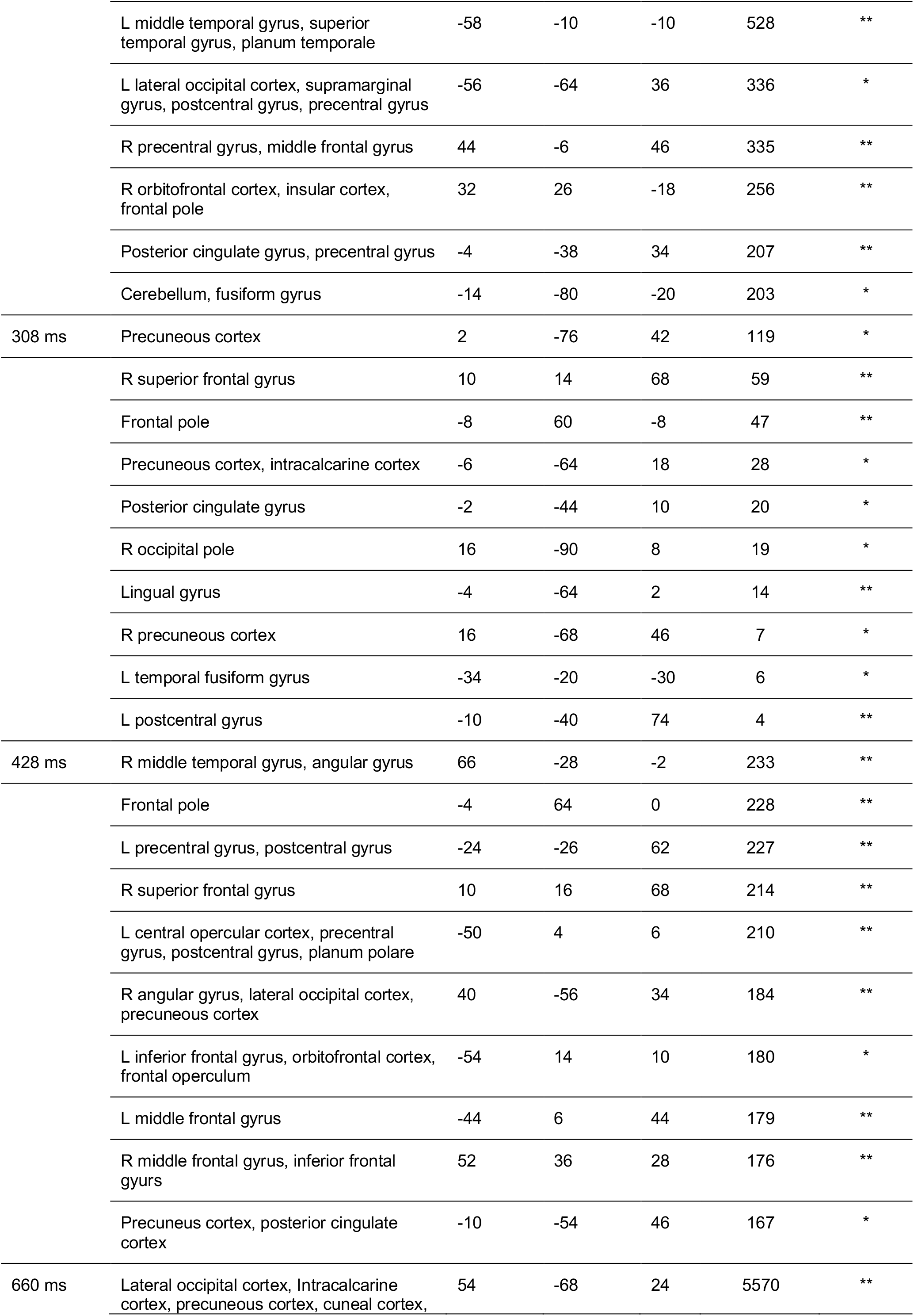
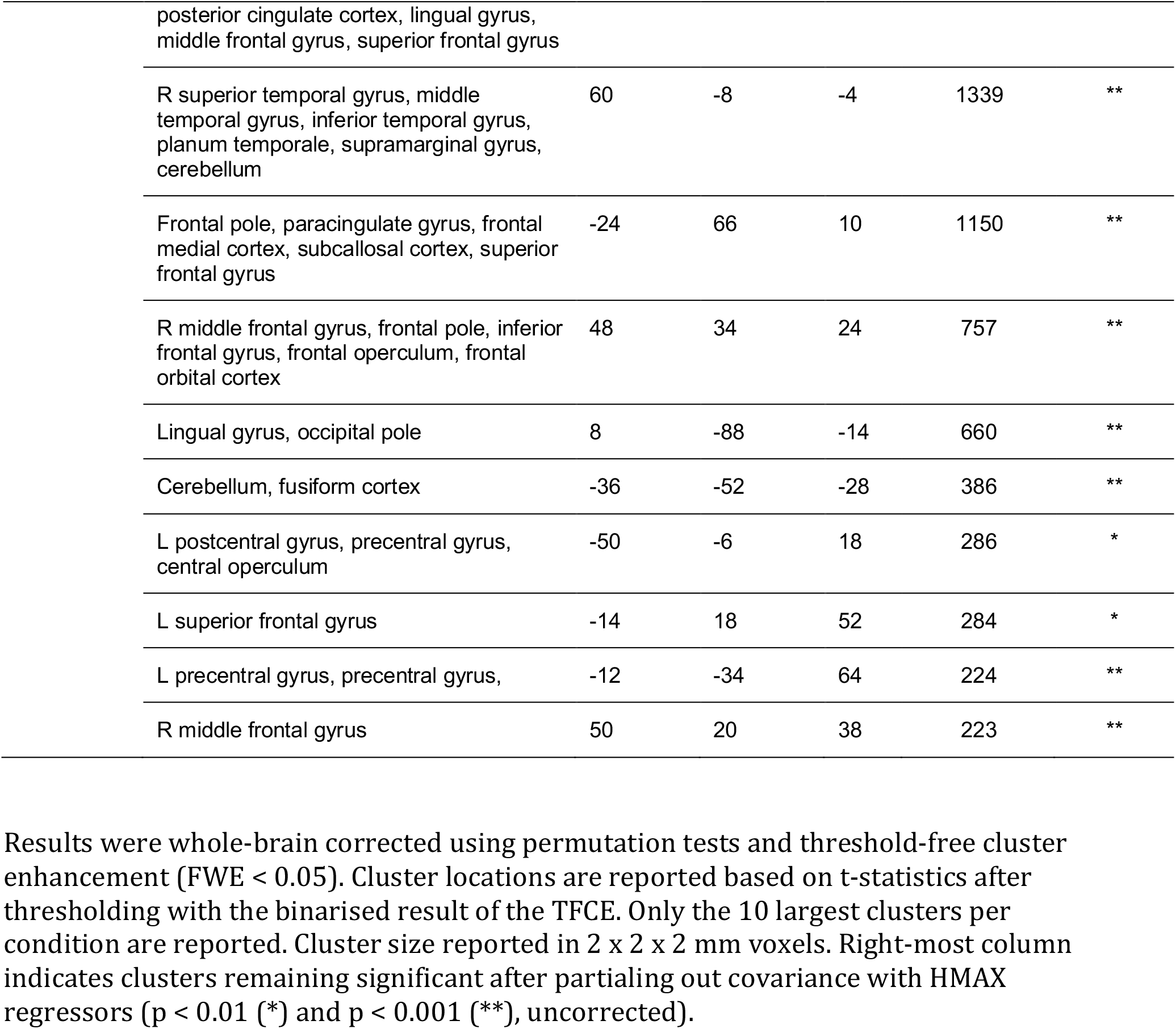
Significant brain regions based on EEG-fMRI RSA for the selected EEG time points

## Discussion

The faces of those closest to us are arguably the most personally relevant social stimuli we encounter and thus might receive processing priority. In this study, we investigated the time course of neural processing of the faces of romantic partners and friends using simultaneous EEG-fMRI and representational similarity analyses. Our results point to the strong and rapid impact of personal relevance on face processing, with shared representations between EEG and fMRI spatial activation patterns in multiple cortical regions starting as early as 100 ms after stimulus onset.

The goal of combining EEG and fMRI data using RSA is to provide a time-resolved measure of brain activity. For this purpose, modality-specific activations are transformed to abstract representations of the stimulus response space, making use of condition-by-condition variability (for review and discussion of limitations, see Cichy and Oliva 2020). By comparing the representational space across modalities, we identified brain regions with corresponding representations to selected time points in the EEG.

These analyses suggest fast modulation of neural processing across a far more widespread collection of cortical regions than predicted. EEG representations which corresponded to personal familiarity were first apparent as early as 108 ms after stimulus onset. Corresponding fMRI representations were found not only in the visual cortex, but also in the ventromedial and medial prefrontal cortex, involved in the monitoring of self-relevant information and value encoding (Northoff et al. 2006; Berridge and Kringelbach 2015). Further early representations were observed in regions linked to multimodal sensory integration and theory of mind (TPJ) (Patel et al. 2019), and episodic and autobiographical memory (posterior cingulate cortex) (Leech and Sharp 2014). Importantly, previous research has identified these regions to be crucially involved in the processing of familiar faces, termed the ‘extended system’ of face processing (e.g., Gobbini et al. 2004; Leibenluft et al. 2004; Ramon et al. 2015; Visconti Di Oleggio Castello et al. 2017; see supplementary information for results of univariate fMRI analyses). The fast and widespread activation of the extended face processing system, which is comprised of brain areas also involved in social cognition and reward encoding serves to highlight the prioritization of genuinely personally relevant information: In contrast to our findings, a previous MEG study using famous faces instead of personally relevant faces reported decoding of face familiarity only after 400 ms (Dobs et al. 2019). However, that study reported increased early representations of gender and identity for familiar compared to unfamiliar faces with latencies of 60 to 100 ms (see also Ambrus et al. 2019). Based on the timeline of their results, the authors speculated that these activations might be based on selective bottom-up amplification of sensory representations. In contrast, although starting somewhat later, our results show that at 100 ms after stimulus onset, corresponding fMRI representations were evident outside of visual areas, including the PFC. This finding is also in line with reports of fast saccadic responses to human faces in general and to personally familiar faces in particular (Crouzet et al. 2010; Visconti di Oleggio Castello and Gobbini 2015).

The EEG-fMRI RSA analyses suggest that personal relevance of faces can be extracted prior to full structural face encoding. These results add to recent EEG and MEG research suggesting that various aspects of faces including identity, emotion, age, and gender, might be represented within 100 ms after stimulus onset (Nemrodov et al. 2016; Vida et al. 2017; Dima et al. 2018). Our results indicate that, apart from visual regions, frontal and midline brain regions might be involved in this process. Based on this circuitry, we suggest that these effects might rely on associative learning of the link between visual stimulus features of partners and friends with their personal relevance and reward value. This explanation is consistent with the findings of MEG/EEG studies reporting fast activations of the PFC in affective associative learning of faces at latencies starting as early as 50 - 80 ms after stimulus onset (for review, see Steinberg et al. 2013), but also with studies pointing to the high importance of anterior regions in the processing of familiar faces (Ramon et al. 2015). Therefore, future models of face perception should consider the influence of face-unspecific mechanisms on face perception, like reward-related processes, which seem to play an important role in processing real-life faces. At 200 ms, shortly after the stage of structural face encoding, significant representations additionally included the fusiform gyrus, but also amygdala, insular cortex and N. accumbens. This time course of representations of face relevance suggests that the response of subcortical relevance- and reward related structures like the amygdala and the N. accumbens might rely on the output of structural face encoding at around 170 ms after stimulus onset. Finally, at the stage of higher-order processing from approximately 400 ms after stimulus onset, significant representations were identified in all regions of the core and extended face processing network, including amygdala, insular and orbitofrontal cortex (Gobbini and Haxby 2007), but also in putamen, cerebellum and regions of the brain stem.

EEG linear regression results suggest that early processing starting around 100 ms after stimulus onset mainly reflects the amplification of familiarity; whereas effects specific for romantic love emerge around 250 ms. For both variables, stable and long-lasting effects are evident at higher-order processing stages. However, it is important to keep in mind that the terms ‘familiarity’ and ‘love’ in this case are not exclusively related to Friend and Partner, respectively, but reflect different aspects of close relationships (Hendrick and Hendrick 1986): Both Partner and Friend are of high personal relevance; and one can feel both friendship love towards a close friend as well as friendship towards one’s partner. Thus, romantic love in our study refers to representations that are unique to a romantic relationship, over and above a close friendship.

In this study, although effects of valence consistent with previous research were present in unimodal fMRI analyses (see supplementary material), EEG regression analyses showed that the representational structure of the EEG signal exclusively reflected conceptual representations of Familiarity and Love. This should not come as a surprise. Being able to rapidly respond to those closest to us is a fundamental ability present from early infancy (Pascalis et al. 1995). It also suggests that emotional expressions are not processed in a rigid, automatic way, but that the representational space of brain responses reflects context-specific relevance of social stimuli.

Some limitations to our study design were imposed by our multimodal setup, limiting the number of trials and stimulus conditions used within a reasonable length session. A larger number of stimulus conditions would provide larger RDMs, able to capture more detailed representational structure, and would reduce the ambiguity in the link between EEG and fMRI (for review, see Cichy and Oliva 2020). Inferring the time course of fMRI using EEG data is based on identification of corresponding representations. Therefore, it is possible that similar representations in different brain regions occur at different time points; however, it has been argued that this is unlikely, given that the transformation of information exchanged between brain regions is complex and non-linear (Cichy and Oliva 2020). More generally, it has also been shown that the size of the stimulus set can impact familiar face processing (Ramon et al. 2019). Expanding the stimulus set poses problems when examining highly personal social relations, particularly romantic partners, though using pictures of faces taken from different visual perspectives is one possibility. In addition, cross-validation of RDMs was not performed, as BOLD response estimates based on half the data were not reliable. However, this problem is mitigated by the multimodal analysis approach; crucially, we do not infer directly on unimodal RDMs, but infer on the similarity between fMRI RDMs and EEG RDMs, and EEG RDMs and model RDMs. Should fMRI or EEG RDMs have spurious, artifact or noise-driven structure, one would not expect robust relationships with model RDMs or RDMs from the other modality. Finally, this study included only young, heterosexual female participants; future studies might examine the extent to which the neural processing of personal relevance is similar or different across different gender identities and sexual preferences. Qualitative differences, however, are presumably most likely across different age groups and durations of relationships.

Understanding the time course and neural circuitry of processing individually relevant faces might be of special importance in clinical conditions, where processing of emotional or social information is specifically disrupted. Autism spectrum conditions (ASC) are an example in which atypical face processing might reflect altered personal relevance attached to strangers, rather than a dysfunction of the neural face processing architecture (Pierce 2004), with potential consequences for the design of targeted interventions.

What is clear is that in basic social neuroscience, as well as in clinical research, there are compelling reasons to study responses to individualised social stimuli. Such studies might yield valuable insights into how the brain engages circuitry beyond that typically considered in models of face perception, in order to prioritise processing of genuinely personally relevant faces. After all, by far the most pervasive social stimuli we encounter in our daily lives, the ones our brains are most attuned to, are our close friends and loved ones.

## Acknowledgements

M.B. and T.J. were supported by the Berlin School of Mind and Brain (Humboldt-Universität zu Berlin) and the Erasmus program. The authors thank Michael Lindner and Catriona Scrivener for assistance with data collection and Luca Brivio for assistance with data analysis.

## Author contributions

M.B., T.J. and I.D. designed the experiment, M.B. and T.J. collected data, conducted analyses and drafted the manuscript, M.B. and O.B. analysed EEG data. All authors provided feedback and revised the manuscript.

## Code and data availability

The code and data is available here:

https://osf.io/rcufv/?view_only=d7fb9219f6da4a32900ab7149d27745a

The authors declare no conflict of interest.

