## Supplementary file for "Rapid neural representations of personally relevant faces"

### Supplementary information

**Table S1: Rating values of experimental stimuli** (means and standard deviations).

| | **Attractiveness** *(1-7)* | **Valence** *(-3 - +3)* | | | **Arousal** *(1-7)* | | |
|---|---|---|---|---|---|---|---|
| | | **Fearful** | **Happy** | **Neutral** | **Fearful** | **Happy** | **Neutral** |
| **Partner** | 6.1 (1.1) | -0.4 (1.5) | 2.3 (0.7) | 0.0 (1.8) | 3.2 (1.7) | 5.4 (1.5) | 3.7 (1.8) |
| **Friend** | 4.1 (1.4) | -1.1 (1.1) | 1.6 (0.8) | -0.6 (1.1) | 2.1 (1.6) | 2.9 (1.9) | 2.5 (1.6) |
| **Stranger** | 4.1 (1.2) | -2.0 (1.1) | 0.5 (1.1) | -0.6 (1.2) | 1.6 (1.1) | 3.2 (2.0) | 2.5 (1.6) |

### Unimodal fMRI analyses and results

Contrasts of interest in unimodal fMRI analyses included effects of Identity and Emotion. Increased activation for Partner > Stranger and Friend > Stranger was observed in right fusiform cortex, precuneus, posterior cingulate cortex, anterior cingulate cortex (ACC), right middle temporal gyrus and posterior superior temporal sulcus (Fig. S1). Additionally, the contrast Partner > Stranger showed activation in the bilateral orbitofrontal cortex, while Friend > Stranger yielded activation in the ventromedial prefrontal cortex. No significant activations were observed in the contrast Partner vs. Friend.
Emotion contrasts yielded activation for Happy > Neutral in the parieto-occipital cortex (including the intracalcarine cortex, lingual gyrus and precuneus), cerebellum and brain stem. Further clusters were located in the medial and ventromedial prefrontal cortex, orbitofrontal cortex, ACC, inferior and superior frontal gyrus, and insular cortex. Subcortically, activation was seen in the left amygdala, bilateral thalamus and left caudate. For Happy > Fear, activation was found in bilateral insula and frontal operculum, and left-lateralized inferior and middle frontal gyrus and lateral occipital cortex. See Table S1 for a complete list of activated brain regions.

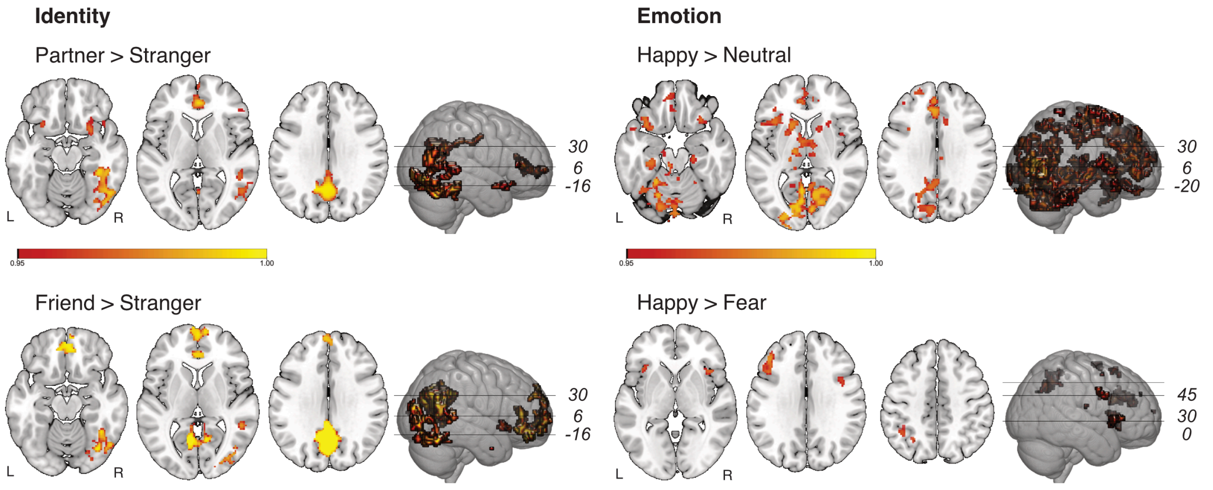

**Fig. S1**. Unimodal fMRI results. Bold activations for Identity comparisons (Partner > Stranger and Friend > Stranger) and Emotion comparisons (happy > neutral, happy > fear). Activation thresholded with FWE < 0.05 using permutations and threshold free cluster enhancement (TFCE).

**Table S2.** Summary of brain activations in unimodal fMRI analyses. Activations are whole-brain corrected using permutation tests and threshold-free cluster enhancement. Clusters were determined from t-statistics after thresholding with the binarised result of the TFCE. Only the 10 largest clusters per condition are reported. Cluster size reported in 2 x 2 x 2 mm voxels.

| **Condition** | **Brain regions** | **Local maxima (mm)** | | | **Cluster size** (voxels) |
|---|---|---|---|---|---|
| | | x | y | z | |
| **fMRI Standard Analyses** | | | | | |
| BF > STR | Fusiform cortex, lateral occipital cortex | 46 | -60 | -22 | 1884 |
| | Precuneous cortex, posterior cingulate gyrus | -4 | -58 | 30 | 1235 |
| | Anterior cingulate cortex, frontal pole, paracingulate gyrus | 0 | 40 | 10 | 495 |
| | R middle temporal gyrus, inferior temporal gyrus, superior temporal sulcus | 50 | -44 | 6 | 201 |
| | R frontal orbitofrontal cortex | 26 | 12 | -16 | 96 |
| | L frontal orbitofrontal cortex | -28 | 18 | -16 | 29 |
| | R Inferior frontal gyrus | 50 | 34 | 8 | 25 |
| | R temporal pole | 44 | 18 | -16 | 16 |
| FR > STR | Precuneous cortex, posterior cingulate cortex, intracalcarine cortex | -2 | -60 | 32 | 2808 |
| | Ventromedial prefrontal cortex, frontal pole, anterior cingulate cortex, paracingulate gyrus, medial superior frontal gyrus | 2 | 50 | -20 | 1571 |
| | R fusiform gyrus, lateral occipital cortex | 44 | -60 | -22 | 1405 |
| | R supramarginal gyrus, middle temporal gyrus, superior temporal sulcus | 50 | -42 | 10 | 98 |
| | Subcallosal cortex | -4 | 20 | -6 | 87 |
| | L lateral occipital cortex | -40 | -92 | 14 | 20 |
| | L lateral occipital cortex | -30 | -88 | 20 | 8 |

| | | | | | |
|---|---|---|---|---|---|
| | L supramarginal gyrus, angular gyrus | -60 | -48 | 12 | 4 |
| | R anterior middle temporal gyrus | 56 | 4 | -32 | 3 |
| Happy > neutral | Lingual gyrus, intracalcarine gyrus, precuneous cortex | -18 | -50 | -14 | 3092 |
| | L Temporal pole, orbitofrontal cortex, insular cortex, frontal operculum, inferior frontal gyrus, caudate, thalamus, pallidum, | -34 | 16 | -32 | 1815 |
| | Frontal pole, anterior cingulate, paracingulate gyrus, superior frontal gyrus | -18 | 68 | 18 | 1665 |
| | Supplementary motor cortex, medial and L superior frontal gyrus, anterior cingulate cortex, mid cingulate cortex | 4 | 6 | 66 | 1109 |
| | Cerebellum, brain stem | 14 | -74 | -38 | 1065 |
| | L precentral gyrus, postcentral gyrus, middle frontal gyrus, | -26 | -16 | 64 | 900 |
| | R cerebellum, brain stem | 32 | -38 | -28 | 424 |
| | L cerebellum | -22 | -82 | -24 | 413 |
| | Cuneal cortex, occipital pole | -8 | -82 | 32 | 288 |
| | Precuneus cortex, posterior cingulate | -6 | -56 | 32 | 254 |
| Happy > fear | L middle frontal gyrus, inferior frontal gyrus, frontal pole | -36 | 42 | 22 | 347 |
| | L Angular gyrus, supramarginal gyrus | -40 | -52 | 46 | 169 |
| | L frontal operculum, insular cortex, | -38 | 22 | 6 | 129 |
| | R insular cortex | 40 | 18 | 0 | 108 |
| | L lateral occipital cortex, superior parietal lobule | -28 | -60 | 42 | 82 |
| | R precentral gyrus, middle frontal gyrus | 42 | 6 | 30 | 67 |
| | L middle frontal gyrus | -34 | 4 | 62 | 49 |
| | L superior frontal gyrus | -20 | 20 | 60 | 44 |
| | L thalamus | -8 | -14 | 18 | 5 |
| | L supplementary motor cortex | -8 | 6 | 54 | 5 |
